# Systemic Sclerosis dermal fibroblast exosomes trigger a Type 1 interferon response in keratinocytes through the TBK/JAK/STAT signalling axis

**DOI:** 10.1101/2023.12.14.570365

**Authors:** Jessica Bryon, Christopher W Wasson, Katja Koeppen, Francesca Chandler, Leon F Willis, Elliott Klein, Elton Zeqiraj, Rebecca L Ross, Francesco Del Galdo

## Abstract

**Background:** Activation of Type I IFN response has been shown to correlates with disease activity in systemic sclerosis. It is currently unknown whether the tissue-specific Type I IFN activation is a consequence of the response observed in blood or rather its source. Exosomes from SSc fibroblasts were recently shown to activate macrophages *in vitro*. Here, we aimed to determine the source of Type I IFN signature in SSc skin biopsies and the potential role of exosomes from SSc dermal fibroblasts in the process.

**Methods:** Skin biopsies were obtained from healthy and SSc patients’ forearms and processed for dermal fibroblasts and keratinocytes. Exosomes were isolated from healthy and SSc dermal fibroblast supernatants by ultracentrifugation and added to human skin keratinocytes. Keratinocyte transcriptome was analysed by RNA-seq analysis. TANK-binding kinase (TBK) and JAK were inhibited using a small molecule inhibitor (GSK8612) and Tofacitinib, respectively.

**Results:** SSc skin biopsies showed highest levels of Type I IFN response in the epidermal layer. RNA-seq analysis of keratinocytes transcriptome following exposure to dermal fibroblast exosomes showed strong upregulation of IFN signature genes induced by SSc exosomes compared to Healthy control. Inhibition of TBK or JAK activity suppressed the upregulation of the IFN signature induced by SSc exosomes.

**Conclusion:** IFN activation of SSc keratinocytes is dependent on their crosstalk with dermal fibroblasts and inducible by extracellular exosomes. Our data indicates that SSc fibroblasts exosomes may carry the ‘‘signal zero’’ of local Type I IFN activation through activation of pattern recognition receptors upstream of TBK.

**Key Messages:**

- SSc patient skin exhibit a type 1 IFN signature with keratinocytes being the major source of the signature
- Cross talk between the fibroblasts and keratinocytes through exosomes may be signal zero for the type 1 IFN signature
- Blocking JAK in the keratinocytes with Tofacitinib disrupts the type 1 IFN signature

## Introduction

Systemic Sclerosis (SSc) is a connective tissue disease driven by tissue and vascular fibrosis of the skin and internal organs. Myofibroblasts have been widely characterised as the key cellular elements of tissue fibrosis for more than two decades. In this context, the key role of pro-fibrotic pathways such as TGF-beta and Wnt has been dissected in fine detail. More recently, the activation of Type I Interferon (IFN) pathway has been associated with markers of disease activity and progression both in skin and lung manifestations but its relationship with fibroblast activation is less clear (1,2,3).

Immunohistochemistry analysis of SSc skin biopsies has revealed that, while there is a clear sign of Type I IFN activation in SSc dermal fibroblasts, the epidermal layer is the major producer of Type I IFN response (2). The trigger for the response in the epidermis is still unclear. Previous studies have shown Cancer Associated Fibroblasts (CAF) exosomes can induce Interferon stimulated genes (ISG) expression in associated epithelial cells (4). In particular, CAF exosomes were shown to trigger Type I IFN activation through uncapped RNA dependent activation of RIG-I in the epithelial cells, which in turn, was dependent on increased activity of RNA Pol III (5). These findings are particularly relevant as SSc fibroblasts share a number of similarities with CAFs (6,7) and autoantibodies against RNA Pol III are one of the SSc specific ANA carrying poor prognostic value for disease progression (8). Consistent with these findings, while our studies were in progress, Bhandari et al have shown that SSc dermal fibroblast exosomes are able to trigger the activation of macrophages *in vitro* (9). These recent data support the previously observed role of dermal fibroblasts in participating to the pathogenesis of SSc beyond tissue fibrosis (10) and possibly in a direct activation of the immune events participating to tissue damage.

In this context, we set out to determine whether SSc dermal fibroblasts could participate in the Type I IFN response observed in SSc skin, particularly through extracellular vesicles. Here we show for the first time that the Type I IFN activation shown by keratinocytes *in vivo* can be driven by SSc fibroblast exosomes. Further, we show that exosomes induced Type I IFN signalling in keratinocytes is dependent upon the exosome RNA induced activation of the TBK1/JAK/STAT signalling cascade.

## Materials and Methods

The Materials and Methods are included as a supplementary file

## Results

### SSc keratinocytes expression of type 1 IFN signature is lost *in vitro* and can be induced by SSc dermal fibroblast co-culture

Previous studies, including our own, have shown that keratinocytes in SSc skin are a major source of ISG expression (2). Consistent with these findings we observed high levels of pSTAT1, CXCL10 and MX1 in the epidermis of SSc skin compared to healthy control biopsies (Figure 1A). We then set out to determine whether SSc keratinocytes could maintain the ISG response when isolated from the skin, similarly to what has been repeatedly observed for the pro-fibrotic activation of dermal fibroblasts (11). SSc keratinocytes isolated from the same biopsies shown in Figure 1A did not display elevated MX1, CXCL10, CXCL11 transcript levels or pSTAT1 protein levels (Figure 1B-E) compared to healthy control keratinocytes, when cultured *in vitro*. Interestingly, these isolated SSc keratinocytes were still able to induce ISG expression upon stimulation with Interferon alpha (Figure 1C-D) suggesting the Type I IFN signalling pathway is still functional in the SSc keratinocytes in culture.

**Figure 1:**
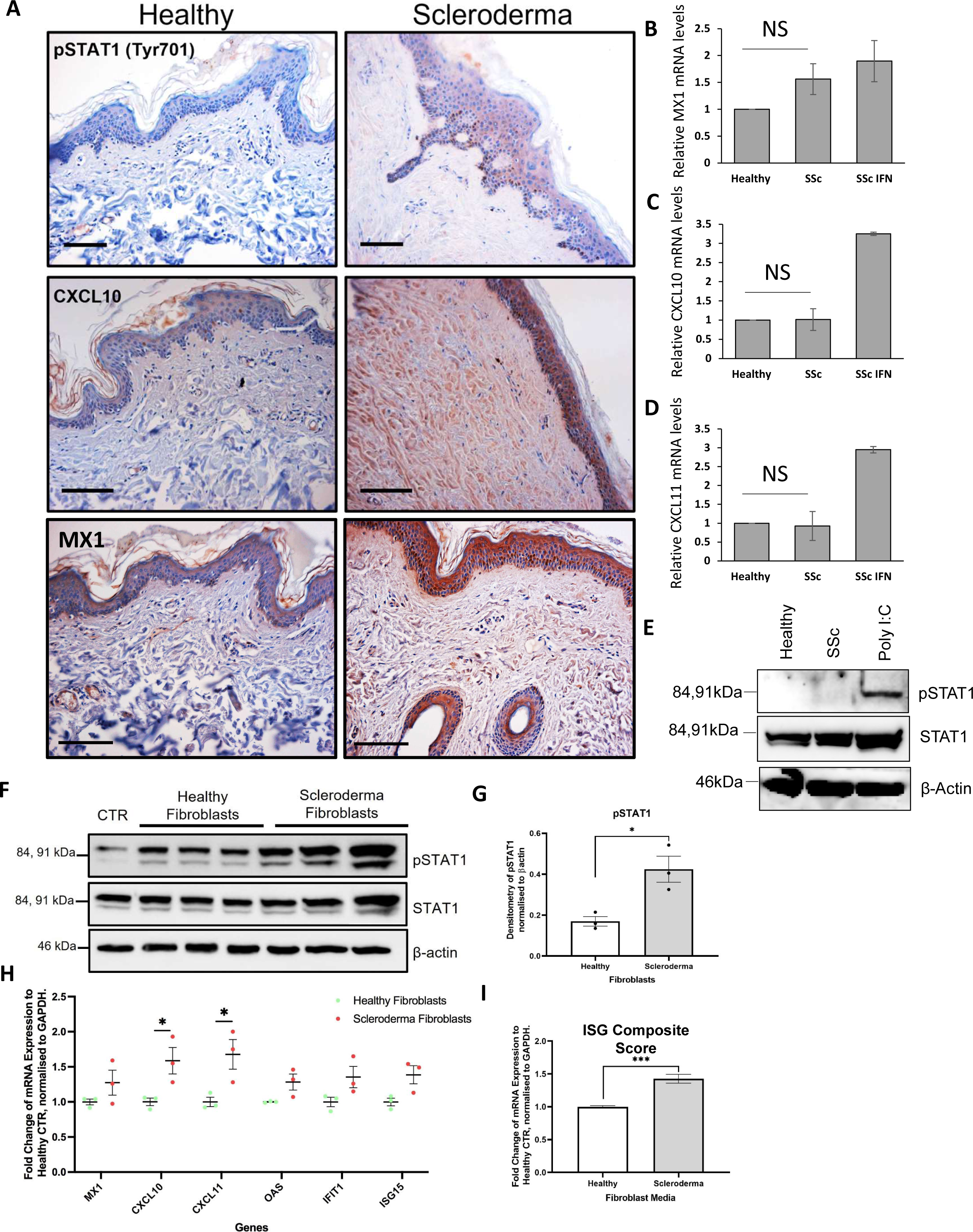
Scleroderma dermal fibroblasts induce type I interferon signalling in keratinocytes. (A) Healthy and SSc skin sections obtained from patient forearms were stained with antibodies specific for phosphorylated STAT1, CXCL10 and MX1. Scale bars represent 50μm. (B-D) SSc skin keratinocytes were isolated from skin biopsies and passaged 3 times; RNA was then isolated. SSc keratinocytes were stimulated with IFN-α (2ng/ml) for 2.5 hours. MX1 (B), CXCL10 (C), CXCL11 (D) transcript levels were assessed by RT-qPCR. Graphs represent fold change releative to healthy control. (E) Protein was isolated from SSc keratinocytes (n=3) and healthy (n=3) control keratinocytes. Lysates from healthy keratinocytes stimulated with POLY I:C (24 hours 10ug/ml) was used as a positive control. pSTAT1 and Total STAT1 protein levels were assessed by western blot. β-actin was used as a loading control. (F-G) Healthy (n=3) and Scleroderma (n=3) fibroblasts were co-cultured in cell culture inserts with human epidermal keratinocytes (HaCats) for 48 hours in serum-free DMEM. (F) pSTAT1 (Tyr701), STAT1 protein levels were assessed by western blot. β-actin was used as a loading control. (G) Protein expression of pSTAT1 was quantified by densitometry and normalised to β-actin levels. (H) Fold change of mRNA transcript expression of ISGs (MX1, CXCL10, CXCL11, OAS, IFIT1 and ISG15) in HaCats co-cultured with healthy fibroblasts, compared to SSc. (I) Fold change of mRNA transcript expression of an ISG composite score of the six genes described in H genes, normalised to GAPDH and ribosomal protein as a house-keeping gene, relative to mean healthy exosome-treated HaCats. *p<0.05. ***?

Next, we assessed whether SSc fibroblasts could trigger Type I IFN response in keratinocytes. To focus our study on the role of the fibroblasts we co-cultured the skin keratinocyte cell line HaCats with 3 different healthy and SSc fibroblasts in a transwell assay (Figure 1F). We observed that both healthy and SSc dermal fibroblasts triggered STAT1 phosphorylation in HaCats compared to the control, with a significantly greater induction in the HaCats cultured with SSc dermal fibroblasts (Figure 1G). Accordingly, SSc dermal fibroblasts induced greater levels of mRNA expression of ISGs (MX1, CXCL10, CXCL11, OAS1, IFIT1 and ISG15) compared to healthy control (Figure 1H and Supplementary Figure 1). Combining the ISG analysed in a composite score as previously described (2), we observed an overall 1.5-fold increase of ISG induction in the HaCats co-cultured with SSc fibroblasts compared to healthy control (Figure 1I).

### Healthy and SSc fibroblast exosomes display similar characteristics

To determine whether fibroblast exosomes could contribute to the induction of Type I IFN in keratinocytes, we isolated exosomes from healthy and SSc fibroblast supernatants. Firstly, we observed that healthy and SSc fibroblasts secreted similar amounts of exosomes as shown by similar levels of exosome marker expression (TSG101 and CD63) in the isolated exosomes (Figure 2A). This was confirmed via CD63 ELISA where we quantified that both healthy and SSc fibroblasts secreted exosomes in the order of 1×10^10^ (from 20 million fibroblasts) (Figure 2B). Further, electron microscopy analysis and dynamic light scattering indicated that the exosomes secreted from healthy and SSc fibroblasts were of similar size and distribution (Figure 2C-D). Finally, we set out to determine the cell entry kinetics of both sets of exosomes as assessed by membrane specific dye staining followed by immunofluorescence analysis. As shown in Figure 2E, we observed no apparent difference in timing and intracellular localisation of fibroblast exosomes from SSc or HC dermal fibroblasts, with a time dependent increase in exosome uptake continuing to rise up to the 48 hours in both sets (Figure 2F).

**Figure 2:**
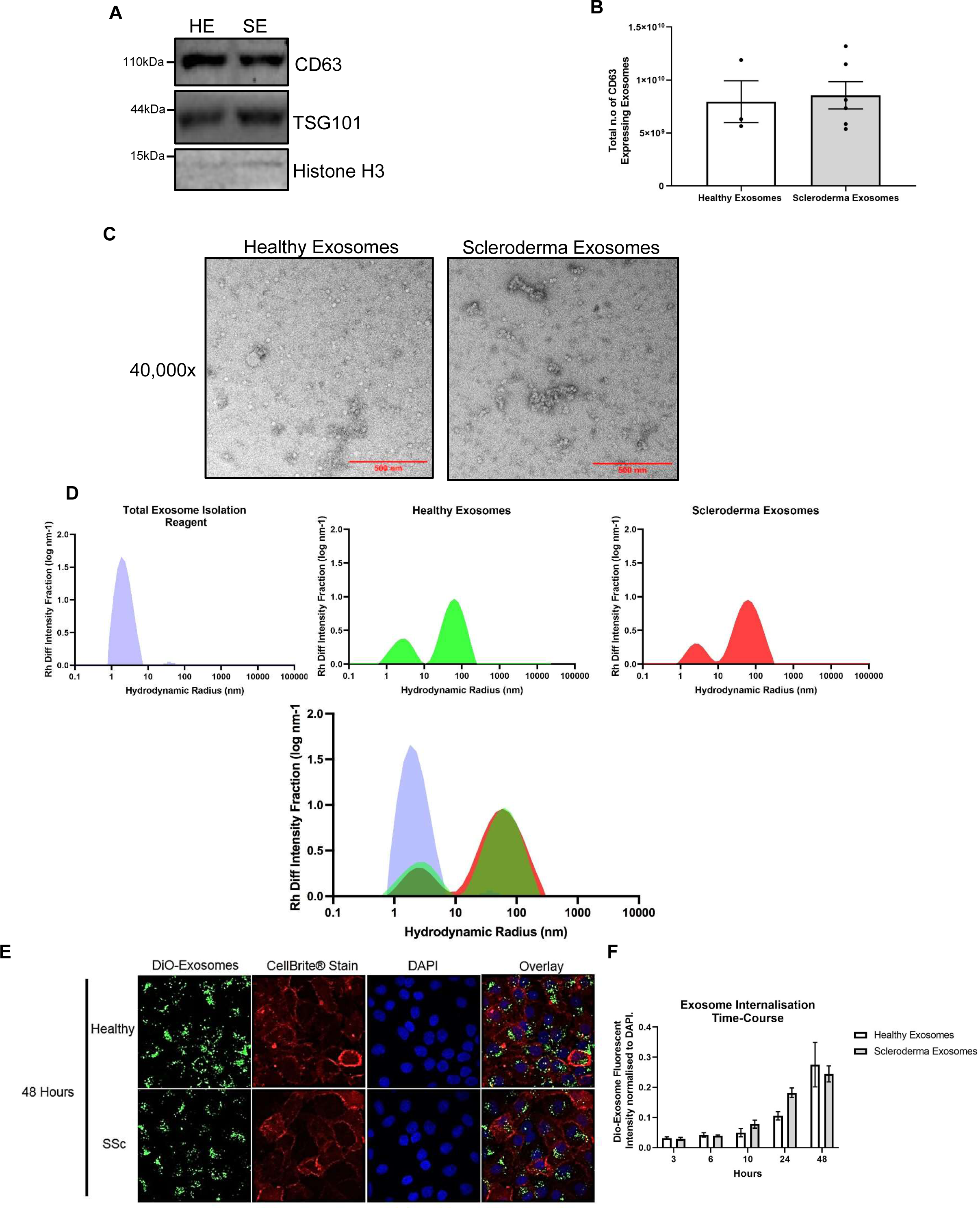
Characterisation of exosomes from healthy SSc dermal fibroblasts. Exosomes were isolated from healthy (n=3) and SSc (n=6) fibroblast media. (A) Isolated exosomes were probed for TSG101, CD63 and Histone H3 by western blot. (B) CD63 ELISA quantification of exosomes from healthy (n=3) and SSc (n=5) patient fibroblasts. (C) Representative transmission electron microscopy images of healthy and scleroderma exosomes at 40,000x magnification. (D) Representive Size distribution histograms of diluted total exosome isolate buffer (TEI), healthy (n=3) and SSc (n=3) exosomes, from regularisation analysis of dynamic light scattering data. (E) Healthy and SSc fibroblast exosomes were stained with DiO-lipid dye and added to HaCats at 3, 6, 10, 24 and 48 hours. (F) Bar charts illustrates the green, fluorescent intensity per image, normalised to include a DAPI intensity, per 3 images over a time-course (3, 6, 10, 24 and 48 hours).

### SSc fibroblast exosomes impart a pro-inflammatory effect on keratinocytes compared to healthy control fibroblast exosomes

To determine the global effects of the SSc fibroblast exosomes on skin keratinocytes, we performed RNA-sequencing (RNA-seq) of HaCats following 48 hours culture in exosome conditioned media (1% total volume) (Figure 3A). Hierarchical clustering of differentially expressed genes (DEGs) showed a clear separation of the transcriptomic response induced by exosomes compared to mock and a separation between SSc and Healthy exosomes (Figure 3B). The RNA-seq analysis revealed there were 1097 DEGs (p < 0.05 & abs (log2 FC) > 1) between HaCats stimulated with healthy fibroblast exosomes compared to control HaCats, 881 DEGs in HaCats stimulated with SSc fibroblast exosomes compared to control HaCats, and 60 DEGs in HaCats stimulated with SSc fibroblast exosomes compared to HaCats stimulated with healthy fibroblasts exosomes. Within the 60 SSc specific DEGs found, 39 were up-regulated (Figure 3C) and 21 were downregulated (Figure 3D) in keratinocytes stimulated with SSc fibroblast exosomes compared to healthy control. The full list of log2 fold changes and p-values from the three comparisons for all genes detected in HaCats can be found in Supplementary File 1.

**Figure 3:**
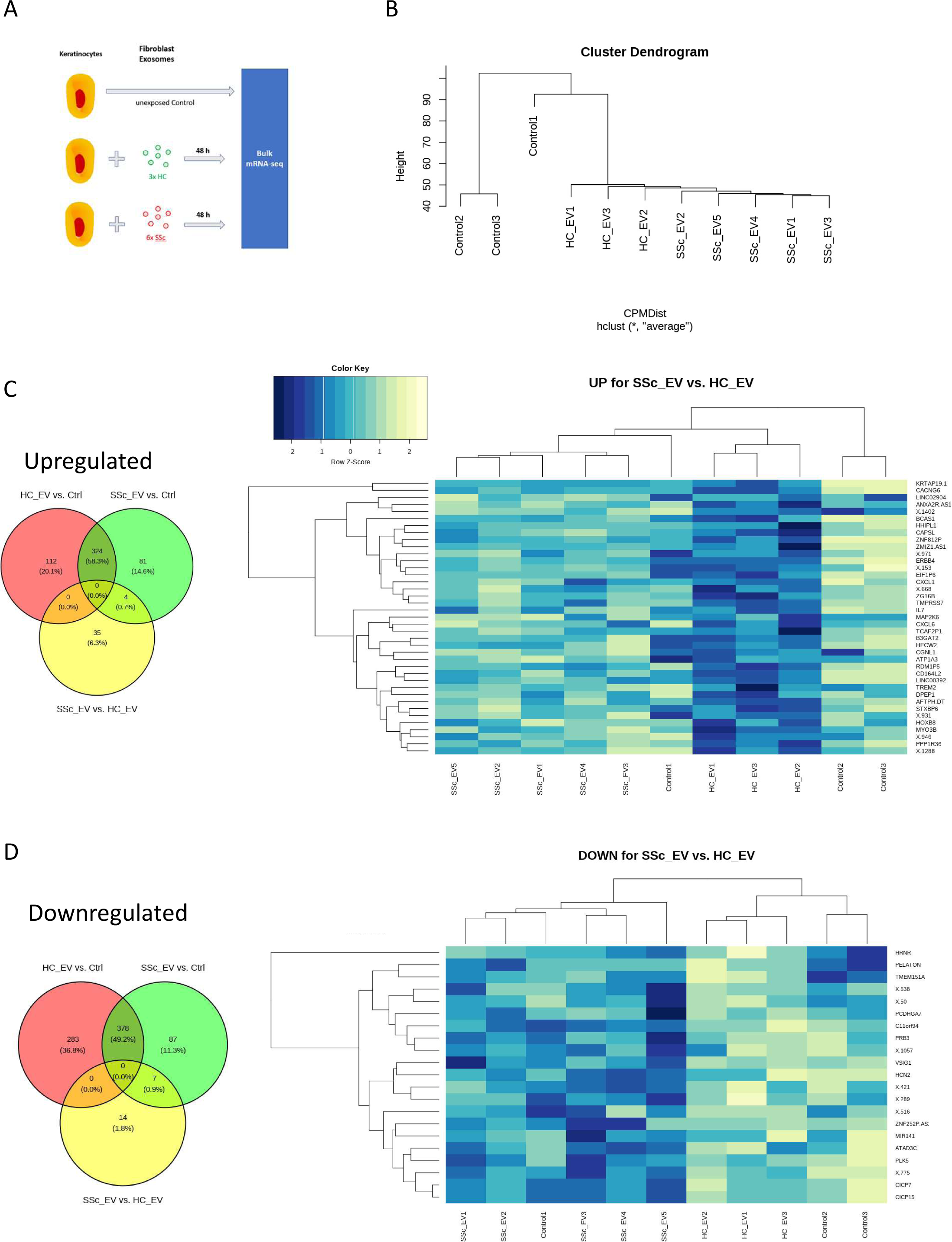
Global gene analysis of Keratinocytes stimulated with healthy and SSc dermal fibroblast exosomes. (A) schematic of experimental approach for the RNA-seq analysis. Hacat keratinocytes were stimulated with healthy (n=3) and SSc (n=5) fibroblast exosomes for 48 hours. RNA was isolated from the stimulated hacats and RNA-seq analysis was performed. (B) Cluster dendrogram clustering of the responses of healthy and SSc in hacats compared to control. Venn diagram and heatmaps of differential expression genes with a fold change greater than 1.5-fold scaled by row for genes upregulated (C) and genes downregulated (D) in Hacats stimulated with SSc fibroblast exosomes compared to healthy fibroblast exosomes. The heat map illustrates significantly (p<0.05) differentially expressed genes (DEGs) and the colour scale represents the row Z-Score, with dark blue representing low gene expression levels and light yellow indicating high expression relative to other samples in the same row.

Gene ontology enrichment analysis using the fold changes for all detected genes revealed that several pathways involved in immune signalling (responses to Type 1 IFN, responses to IFN gamma, humoral immune responses and response to chemokines) were upregulated in HaCats stimulated with SSc fibroblast exosomes compared to healthy fibroblast exosomes (Figure 4A). Analysis of individual genes showed a number of ISGs were differentially upregulated in HaCats stimulated with SSc fibroblast exosomes, as shown in the volcano plot (Figure 4B). A list of the 25 most significant DEG are shown in Figure 4C. The list includes a number of ISGs. Further analysis of Type I IFN signalling in HaCats stimulated with healthy and SSc dermal fibroblast exosomes compared to the mock control revealed that the healthy dermal fibroblast exosomes impart an anti-inflammatory response on the keratinocytes as shown by reduced expression of a number of ISGs. Expression levels of IL-7, CXCL6, CXCL1, GBP1, MMP7, TREM2, and CTSS were significantly reduced in HaCats stimulated with healthy fibroblast exosomes compared to unexposed HaCats (Figure 4D). Expression levels of these ISGs were also significantly reduced in HaCats stimulated with healthy control fibroblasts compared to HaCats stimulated with SSc fibroblasts and not significantly different between HaCats stimulated with SSc dermal fibroblast exosomes and unexposed control HaCats (Supplemental Figure 2). Collectively, these data suggest that an important function of exosomes in healthy skin is to dampen pro-inflammatory IFN signalling in keratinocytes, thus contributing to homeostasis, and that the lack of this anti-inflammatory effect of exosomes secreted by SSc fibroblasts on keratinocytes may lead to loss of homeostasis and contribute to a more pro-inflammatory state in SSc patients.

**Figure 4:**
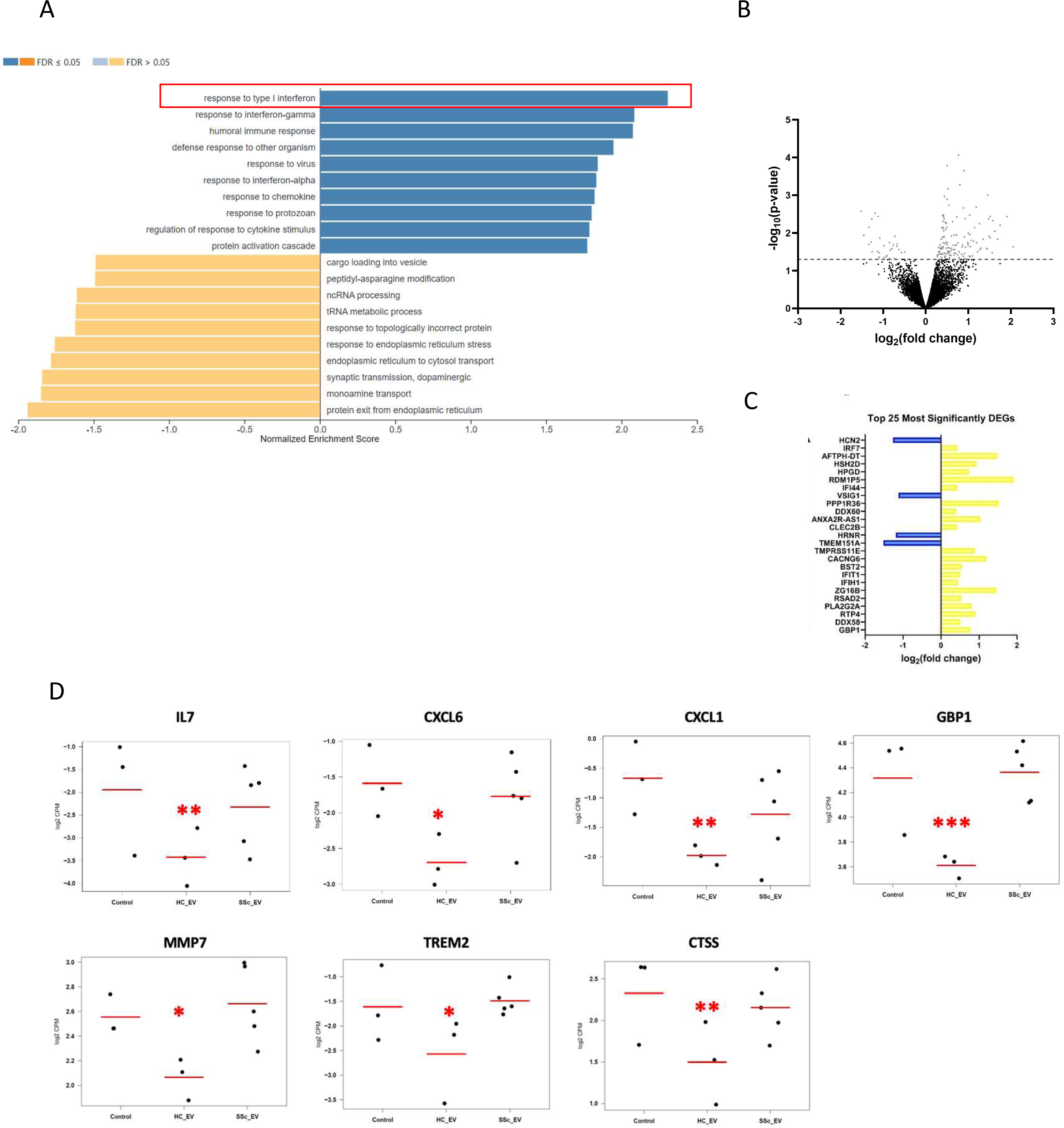
SSc fibroblast exosomes impart a pro-inflammatory effect on keratinocytes compared to Healthy control fibroblast exosomes. (A) Gene ontology (GO) enrichment analysis of biological processes associated with DEGs, in HaCats treated with SSc exosomes compared to healthy control exosomes. Dark blue or orange = false discovery rate (FDR) < 0.05; light blue or orange FDR > 0.05. (B) Volcano plot of log2 fold change against -log10 p-value for each DEG in HaCats stimulated with SSc fibroblast-derived exosomes, when compared against healthy exosomes. Red = significant DEGs. Blue = significant DEGs associated with interferon signalling. (C) A list of the 25 most significant DEG from the analysis. (D) Gene expression levels of selected ISGs (IL-7, CXCL6, CXCL1, GBP1, MMP7, TREM2 and CTSS). Comparisons made between control Hacats and Hacats stimulated with healthy and SSc dermal fibroblast exosomes. *P < 0.05, **P < 0.01, *** P < 0.001

Previous studies have shown SSc fibroblast (9) and CAF exosomes (4,5) actively stimulate macrophages and epithelial cells compared to healthy control fibroblast exosomes. Here we show healthy dermal fibroblast exosomes negatively regulate Type I IFN signalling and this ability is lost in SSc fibroblasts, causing an increase Type I IFN signature in HaCats relative to HC exosomes. To validate this finding, we performed a qRT-PCR based superarray to assess 79 Type I IFN-dependent ISGs on RNA isolated from HaCats treated with healthy and SSc dermal fibroblast exosomes. Volcano plot analysis on three independent experiments showed 16 ISGs upregulated by SSc exosomes compared to Healthy exosomes (Figure 5A-B and Supplementary Figure 3). Altogether, the qRT-PCR based IFN array validated the RNA-seq data indicating a strong Type I IFN response in HaCats stimulated with SSc fibroblast exosomes compared to healthy control exosomes. At the protein level, whole cell protein lysates from the HaCats in the same experimental conditions showed a time dependent increase in pSTAT1 levels (Figure 5C-D), consistent with the RNA findings (Figure 5A-B) and the kinetics of exosome internalisation (Figure 2E). Accordingly, the ISG composite score showed a similar increment over time becoming significantly induced by SSc exosomes at 24 and 48 hours (Figure 5E).

**Figure 5:**
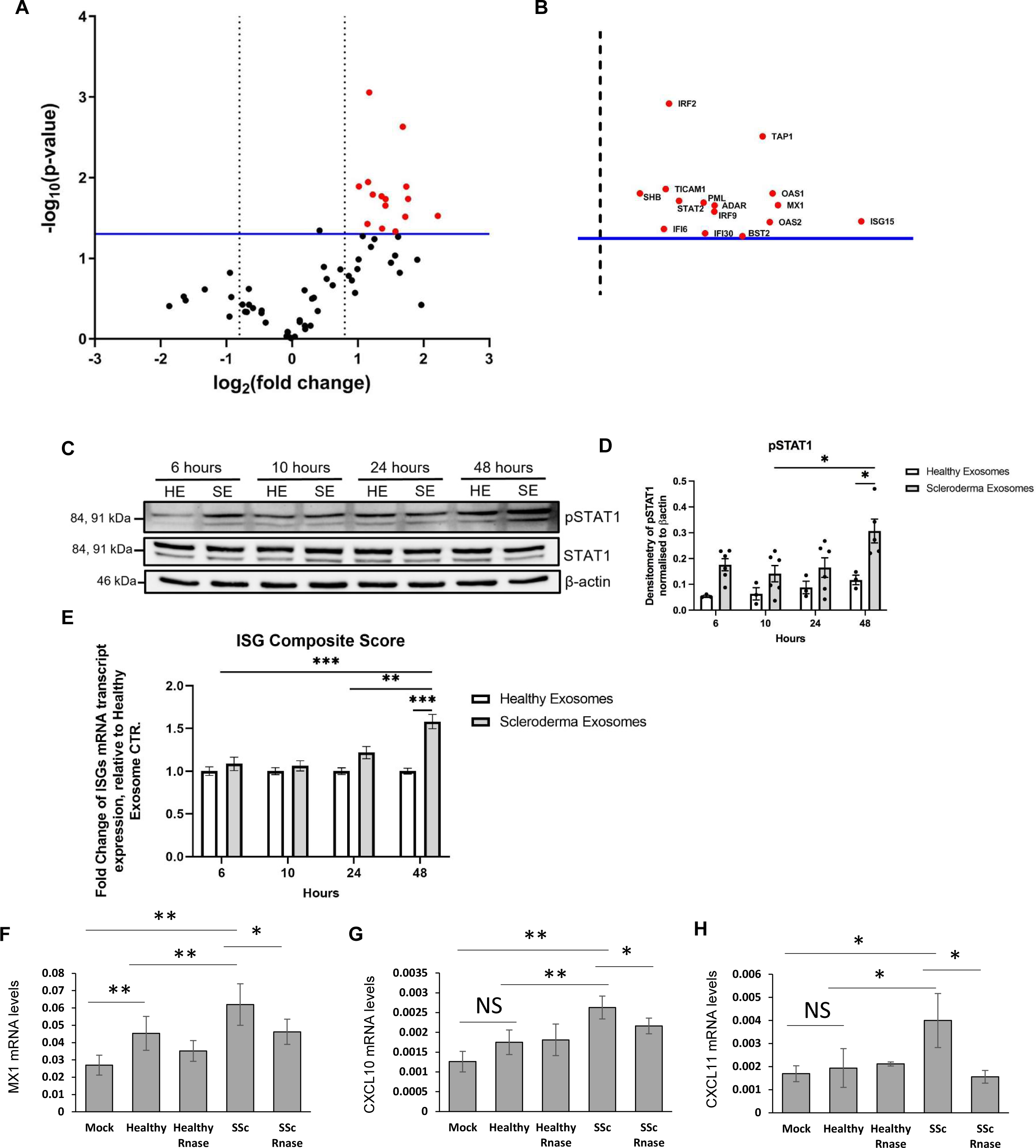
Scleroderma dermal fibroblast exosomes induce Type 1 IFN signalling in keratinocytes through their RNA content. HaCat keratinocytes were stimulated with healthy and SSc fibroblast exosomes for 48 hours. RNA was extracted from the stimulated HaCats. A Type 1 IFN superarray was performed (A) Volcano plot which shows the fold change of gene expression of interferon-type-1-stimulated genes in HaCats stimulated with SSc exosomes relative to HaCats stimulated with healthy exosomes. (B) An enlarged version of the volcano plot in illustrating the IDS of the genes significantly upregulated in SSc vs healthy exosome treated HaCats. HaCat keratinocytes were stimulated with healthy and SSc fibroblast exosomes for 6-48 hours. Protein and RNA were isolated from stimulated HaCats. (C) pSTAT1 (Tyr701) and STAT1 protein levels were assessed by western blot. β-actin was used as a loading control. (D) Bar chart illustrates quantification of protein expression by densitometry and normalised to β-actin levels. Data represents means ± SE. Data was checked for normality and statistically analysed using a Two-way multiple comparison ANOVA, normalised by Tukey. (E) Bar chart illustrates the fold change of mRNA transcript expression of an ISG composite score of multiple interferon-related genes normalised to GAPDH, relative to healthy exosome stimulations, over a time course. ISG composite score included 6 ISGs; MX1, CXCL10, CXCL11, OAS, IFIT1 and ISG1. Data represents means ± SE. Data was checked for normality and statistically analysed using a Two-way multiple comparison ANOVA, normalised by Tukey. RNA was extracted from healthy and SSc fibroblast exosomes and transfected into HaCat cells for 48 hours. In addition the RNA was pre-treated with RNase prior to transfection. MX1 (F), CXCL10 (G) and CXCL11 (H) transcript levels were assessed by RT-qPCR. *p < 0.05, **p < 0.01, ***p < 0.001, ****p < 0.0001.

### SSc fibroblast exosomes trigger Type 1 IFN signalling in keratinocytes through a TBK1/JAK dependent response to their RNA cargo

Previous studies have shown that specific uncapped RNA transcripts within CAFs exosomes can trigger Type I IFN signalling in epithelial cells (7). Therefore, we wanted to determine if the RNA contained within the SSc fibroblast exosomes contributed to the Type I IFN stimulation observed in keratinocytes. RNA was extracted from healthy and SSc dermal fibroblast exosomes and transfected into HaCat cells (50ng RNA). Healthy fibroblast exosome RNA had minimal effects on MX1, CXCL10 and CXCL11 expression in HaCats compared to control cells. SSc fibroblast exosome RNA induced 2.5-fold increase in MX1 (p=0.002); 2-fold increase in CXCL10 (p=0.0015) and 3-fold increase in CXCL11 (p=0.016) expression in HaCats (Figure 5F-H). Pre-treatment of exosome extracts with RNase significantly suppressed the upregulation of MX1, CXCL10 and CXCL11, although with variable efficiency (Figure 5F-H). This suggests that the RNA within the exosomes directly contributed to Type I IFN response of HACAT, *in vitro*.

There is a number of ways RNA cargo could induce a Type I IFN response, such as a miRNA induced upregulation of Type I IFN or a direct stimulation of RNA sensors, as it has been described for CAF derived exosomes (4–5). Most RNA pattern recognition receptors (TLR-7, RIG-I and MDA5) signal through TBK1. Therefore, we used the specific TBK1 inhibitor GSK8612 to interrogate the pathway in response to exosome stimulation. The observed increased STAT1 activation by SSc fibroblast exosomes was completely abolished with the TBK1 inhibitor (Figure 6A-B). Accordingly, TBK1 inhibition also suppressed the induction of ISG expression in the HaCats (Figure 6C). Taken together this suggests the SSc fibroblast exosomes induce Type I IFN signalling in HaCats through TBK1. TBK1 is known to activate STAT1 through IRF3/7 in response to viral infections (12). Interestingly the SSc fibroblast exosomes did not activate IRF3 in keratinocytes (Supplementary Figure 4). This suggests that TBK1 activates STAT1 through an alternative pathway. This was further highlighted in the type 1 IFN array and RNA-seq data where IFN alpha/beta expression was not induced by the SSc fibroblast exosomes.

**Figure 6:**
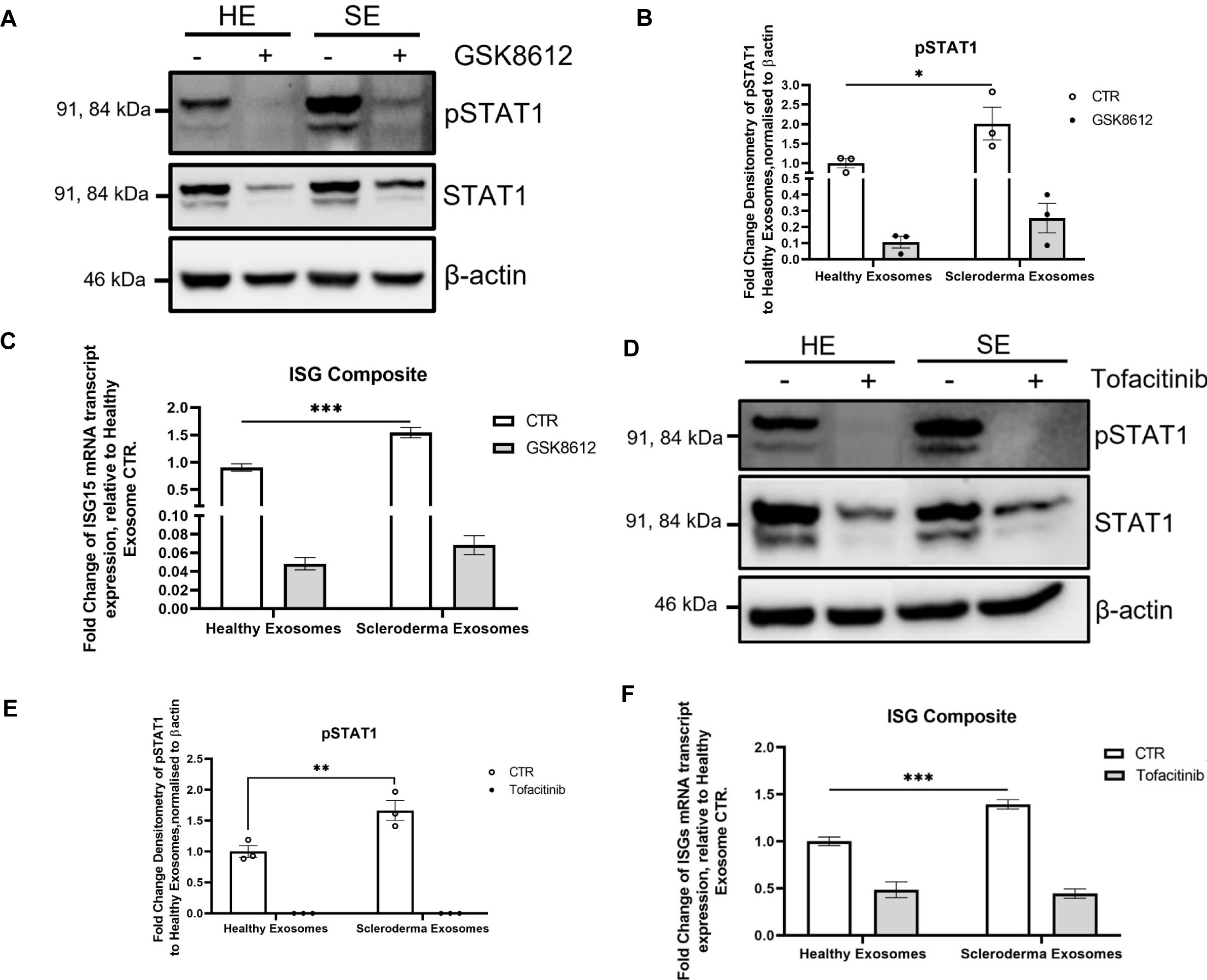
Inhibition of Tank-binding Kinase and JAK-STAT blocks SSc fibroblast exosome mediated Type I IFN signalling in HaCats. (A) HaCats were stimulated with healthy and SSc fibroblast exosomes for 48 hours. In addition the HaCats were treated with 10 µM GSK8612, 1 hour prior to stimulation. pSTAT1 and STAT1 protein levels were assessed by western blot. β-actin was used as a loading control. (B) The bar chart illustrates densitometry analysis of pSTAT1. Data represents means ± SE from three biological repeats. (C) Fold change of mRNA transcript expression of multiple interferon-related genes, normalised by housing keeping gene GAPDH, and relative to untreated healthy exosome-treated HaCats. Data represents means ± SE from three biological repeats (healthy), and four biological repeats (SSc). ISG composite score included 6 ISGs; MX1, CXCL10, CXCL11, OAS, IFIT1 and ISG15. Statistical analysis was performed using a multiple comparison two-way ANOVA, normalised to Tukey. **p<0.01. ***p<0.001. (D) HaCats were stimulation with healthy and SSc exosomes for 48 hours. In addition the HaCats were treated with 20 µM Tofacitinib 1 hour prior to stimulation. pSTAT1 and STAT1 protein levels were assessed by western blot. β-actin was used as a loading control. (E) The bar chart illustrates densitometry analysis of pSTAT1. Data represents means ± SE from three biological repeats. (F) Fold change of mRNA transcript expression of multiple interferon-related genes, normalised by housing keeping gene GAPDH, and relative to untreated healthy exosome-treated HaCats. Data represents means ± SE. ISG composite score included 6 ISGs; MX1, CXCL10, CXCL11, OAS, IFIT1 and ISG15. Statistical analysis was performed using a multiple comparison two-way ANOVA, normalised to Tukey. **p<0.01. ***p<0.001.

To determine whether the TBK1 induced activation of STAT was JAK dependent we used the JAK1 inhibitor Tofacitinib in the same experimental conditions of Figure 6A-B. Similar to TBK inhibition, Tofacitinib completely blocked the SSc fibroblast exosome mediated STAT1 activation in HaCats (Figure 6D-E) as well as ISG expression (Figure 6F), suggesting that the observed Type I IFN response is mediated by activation of JAK. Taken together this suggests the SSc fibroblast exosomes induce a Type I IFN response in keratinocytes through a TBK1-JAK signalling axis (Figure 7)

**Figure 7:**
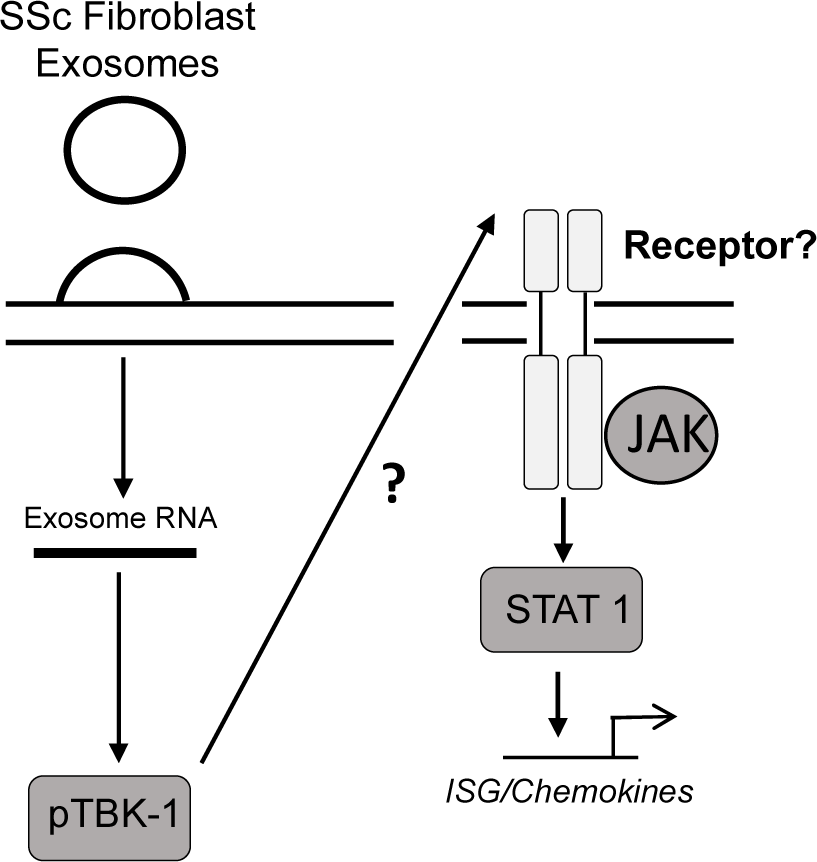
Systemic Sclerosis dermal fibroblast exosomes trigger a Type 1 interferon response in keratinocytes through the TBK/JAK/STAT signalling pathway.

## Discussion

In this study we have shown the Type I IFN response observed in SSc skin is mainly driven by keratinocytes. Importantly we also show that keratinocytes are “responding” to a primary trigger in their microenvironment as they lose any sign of Type I IFN activation once cultured *in vitro*. Parallel to this evidence, co-culture experiments clearly indicate that SSc dermal fibroblasts are able to induce Type I IFN activation in keratinocytes, which can be induced by isolated exosomes. These data suggests that dermal fibroblasts may participate to the pathogenesis of SSc not only for their pro-fibrotic activation but also triggering and/or sustaining the Type I IFN response observed in the tissue. Further, these data also suggests that the increase Type I IFN-inducible chemokines observed in peripheral blood may be a consequence of a tissue driven Type I IFN response rather than its source. Similar data have been suggested in the pathogenesis of SLE (13). This interpretation would also be consistent with the observation that serum proteome correlated with skin transcriptome better than peripheral blood transcriptome in SSc (14). The transfection of exosome cargo RNA in human keratinocytes suggested that the RNA contained within SSc exosomes is contributing, if not driving the Type I IFN response Finally, we interrogated the signalling pathway behind this response with small molecule inhibitors. Although a fine dissection of the pathway(s) fell outside the scope of this study, we were able to demonstrate that SSc fibroblast exosomes induce ISG expression through a TBK1/JAK/STAT1 signalling cascade.

We believe that these data contribute to identifying the source of Type I IFN activation in SSc skin, but we also acknowledge that there are still a number of unanswered questions regarding this phenomenon. Specifically, we still do not have a clear understanding of the factors downstream of TBK1. One standout candidate is IRF3/7. TBK1 has been shown to phosphorylate and activate IRF3/7 in response to viral and bacterial infections which in turn drives a Type I Interferon response (12). Interestingly we have shown that SSc fibroblasts exosomes do not activate IRF3 (Supplementary Figure 4). This suggests TBK1 induces ISG expression through alternative downstream targets. One possible target is NFκB which is known to be a downstream target of TBK1 (15). This hypothesis will be explored in future studies.

The specific exosome cargo that triggers the Type I IFN response in the keratinocytes is not fully understood. We have shown that exosome RNA can induce ISG expression in the keratinocytes but this does not rule out the possibility that proteins or other second messengers in the cargo could also play a role. The exosomes may contain cytokines, which upon exosome uptake could trigger the signalling cascade we have described in the results. With regards to the RNA cargo, the exosomes could contain unshielded/uncapped RNA as described in the CAF work (4,5). In this regard, the uncapped RNA (RN7SL1) has been shown to be directly linked to Notch/c-Myc/RNA Pol III function. We have shown Notch signalling is enhanced in SSc (16,17) and RNA Pol III has been implicated previously (18). This suggests the exosome RNA from SSc fibroblasts may lead to the activation of the Type I IFN pathway in the keratinocytes through recognition of uncapped RNA structures. Therefore, future work will focus on proteomic and RNAseq analysis of SSc fibroblast exosome cargo to determine if cytokines or the specific RNA transcript RN7SL1 are present in the exosomes. Exosomes are known carriers of miRNAs and a number of miRNA are known to regulate Type I IFN responses. Furthermore a number of differentially expressed long non-coding RNAs in SSc have been implicated in regulating interferon responses (19). Therefore, it is possible that a number of miRNA or lncRNA within the exosomes may regulate the responses we have observed in the keratinocytes.

Interestingly, the RNA-seq data revealed that healthy fibroblast exosomes have an anti-inflammatory effect on HaCats that is lost in SSc fibroblast exosomes have a counteractive effect. This sheds a new light on the possible role of healthy fibroblasts exosomes in regulating the immune response to maintain skin homeostasis, whereas this homeostatic regulation is disrupted with SSc fibroblast exosomes. When healthy fibroblast exosomal RNA was isolated and transfected into keratinocytes the RNA did not have an anti-inflammatory effect (Figure 5F-H) but SSc fibroblast exosome RNA did trigger a Type I IFN-ISG response. This suggests proteins within the healthy fibroblast exosomes may impart the anti-inflammatory response but the RNA from SSc fibroblast exosomes reverses this effect.

The ability of the JAK inhibitor Tofacitnib to prevent SSc fibroblast exosome mediated ISG expression is intriguing. These data suggest exosome mediated ISG expression is triggered by cytokine receptor mediated activation of STAT1. The precise cytokine receptor involved is still unknown (Figure 6). There are a number of receptors that may play a role including IL-6 and IFN-1 receptor. Future work will involve testing the supernatants of exosome stimulated keratinocytes for candidate stimulatory cytokines.

An exciting element of the Tofacitnib data is that this drug is approved in the clinic for rheumatoid and psoriatic arthritis. Early clinical trials in SSc have had promising outcomes (20). Tofacitinib blocked ISG signatures in both fibroblasts and keratinocytes. Our data follow that produced in the trial and adds an extra layer of mechanistic data by suggesting the compound is blocking the effects of the ISG signature driven through the crosstalk between these cells.

Bhandari *et al* showed that SSc fibroblast exosomes can induce macrophage activation (9). This observation is complementary to our data showing the same exosomes can induce a Type 1 Interferon response in keratinocytes. It remains to be determined what factor/s within the exosomes lead to the activation of the macrophages and keratinocytes. Therefore, it would be interesting to investigate if the RNA contained within the exosomes can induce the response in macrophages. In addition, it would be interesting to investigate if the supernatant from the exosome stimulated macrophages could induce a response in the keratinocytes and vice-versa. This could result in an immune signalling cascade in patient skin with the fibroblast exosomes as the culprit of this cascade. Bhandari *et al* showed the supernatant from exosome stimulated macrophages could induce myofibroblast activation in naïve fibroblasts. This suggests the exosomes induce a potent response in the macrophages. Unfortunately, from the study we were unable to conclude if the healthy dermal fibroblast exosomes imparted an anti-inflammatory response in the macrophages (similar to the keratinocytes) as the authors did not show a comparison between unstimulated macrophages and macrophages stimulated with healthy fibroblast exosomes.

In conclusion, we have proposed a mechanism behind the Type 1 IFN response found is SSc skin keratinocytes and our experimental data suggests that tissue fibroblasts may be at least one of the sources of Type I IFN activation observed in SSc.

## Supporting information

Supplementary Dataset for DEG genes

## Acknowledgements

This work was funded by a Boehringer Ingelheim opn2expert funding call grant to CWW and FDG. JB was funded by a NIHR BRC PhD studentship (Scleroderma workstream) to FDG. FDG is partially supported by NIHR BRC. EZ was supported by a Wellcome Senior Research Fellowship (222531/Z/21/Z). FC was supported by a Wellcome Trust PhD studentship (219997/Z/19/Z). We thank Prof Sheena Radford (Leeds) for access to the DLS machine, funded by the Medical Research Council (G0900958).

## Contribution

Conceptualisation: C.W.W, R.L.R, E.K, F.D.G

Funding acquisition: C.W.W, F.D.G

Conducting Experiments: J.B, C.W.W, R.L.R

Acquiring Data: J.B, C.W.W, F.C

Analysing Data: J.B, C.W.W, K.K, F.C, L.F.W, E.K, E.Z, R.L.R, F.D.G.

Writing Manuscript: J.B, C.W.W, K.K, R.L.R, F.D.G

All authors reviewed and approved the final version of the manuscript

## Figure Legends

**Supplementary Figure 1:**
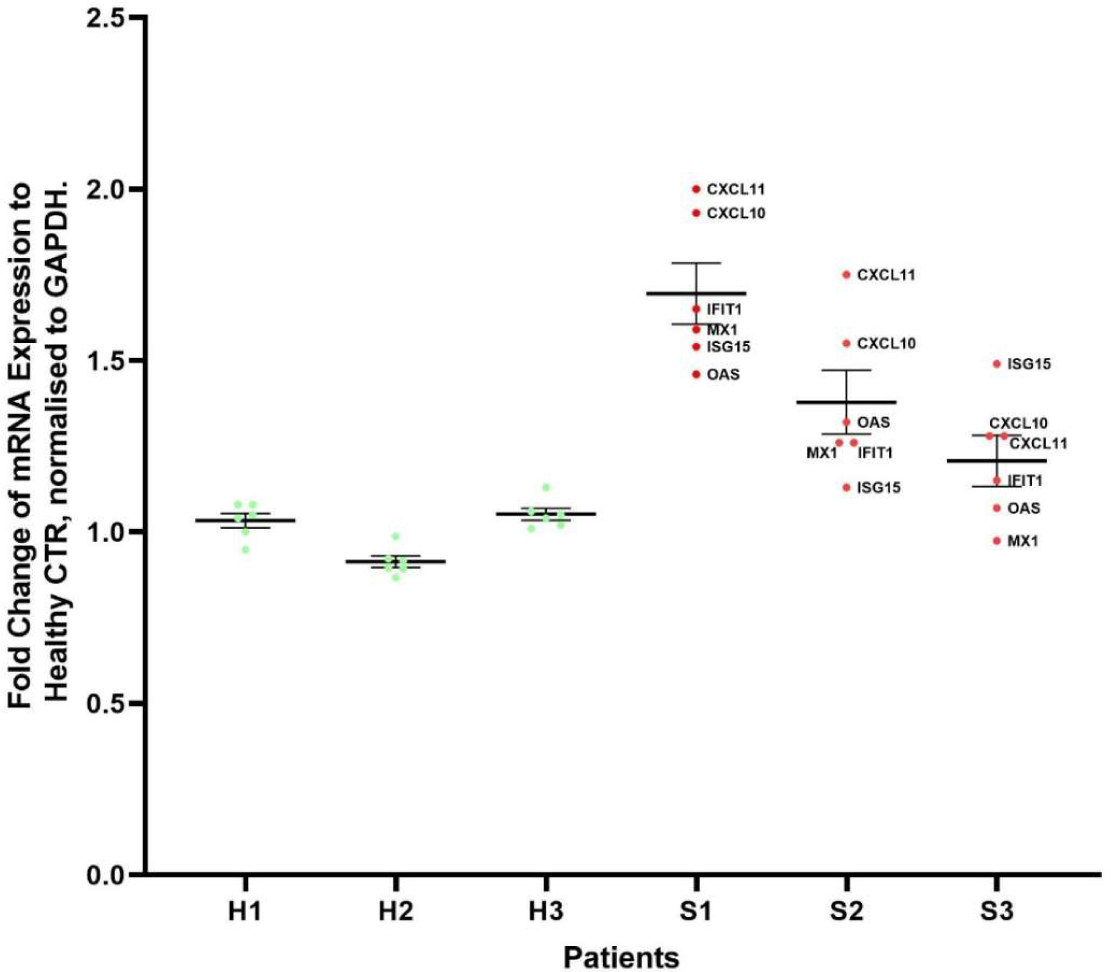
Scleroderma dermal fibroblasts induce type I interferon signalling in keratinocytes. Average ISG transcript expression in keratinocytes co-cultured with each individual SSc patient fibroblast cell line, compared to healthy fibroblast co-cultured HaCats. Data represents means ± SE from three biological repeats. Statistical analysis involved a two-tailed student t-test, assuming equal variance. *p<0.05. ***p<0.001.

**Supplementary Figure 2:**
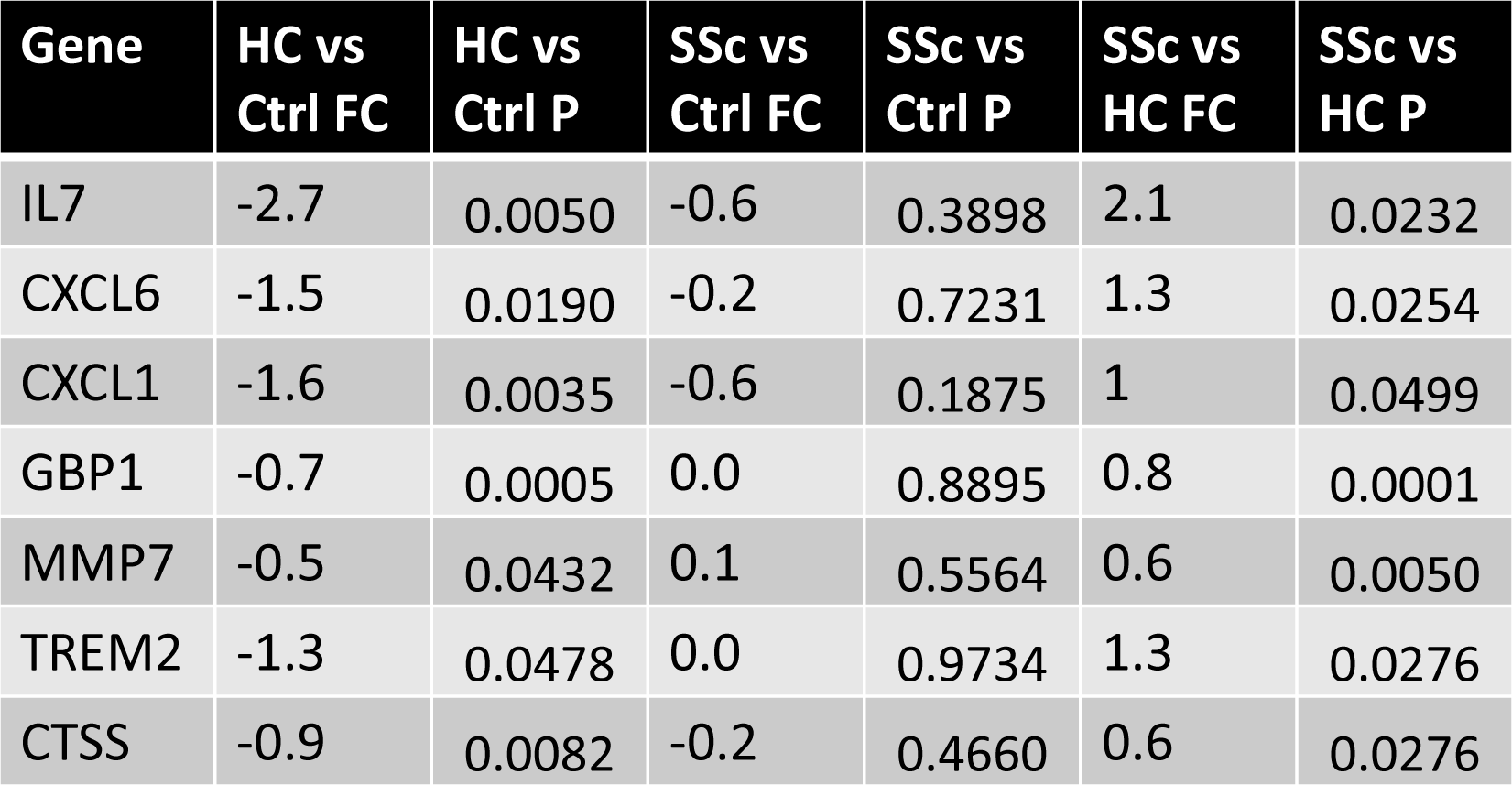
SSc fibroblast exosomes impart a pro-inflammatory effect on keratinocytes compared to Healthy control fibroblast exosomes. Table represents fold change of each inflammatory genes highlighted from the RNA seq analysis.

**Supplementary Figure 3:**
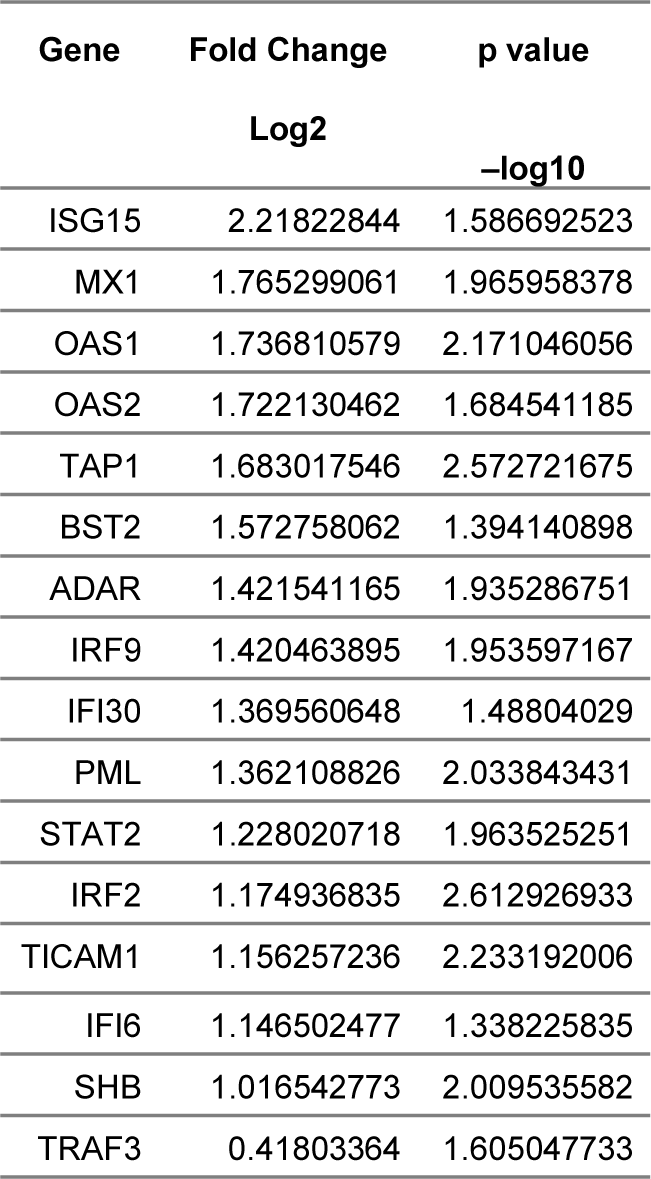
SSc fibroblast-derived exosomes induce interferon-stimulated genes and pSTAT1 in keratinocytes, after 48 hours. Table represents the log fold change of each ISG included in the type 1 IFN superarray.

**Supplementary Figure 4:**
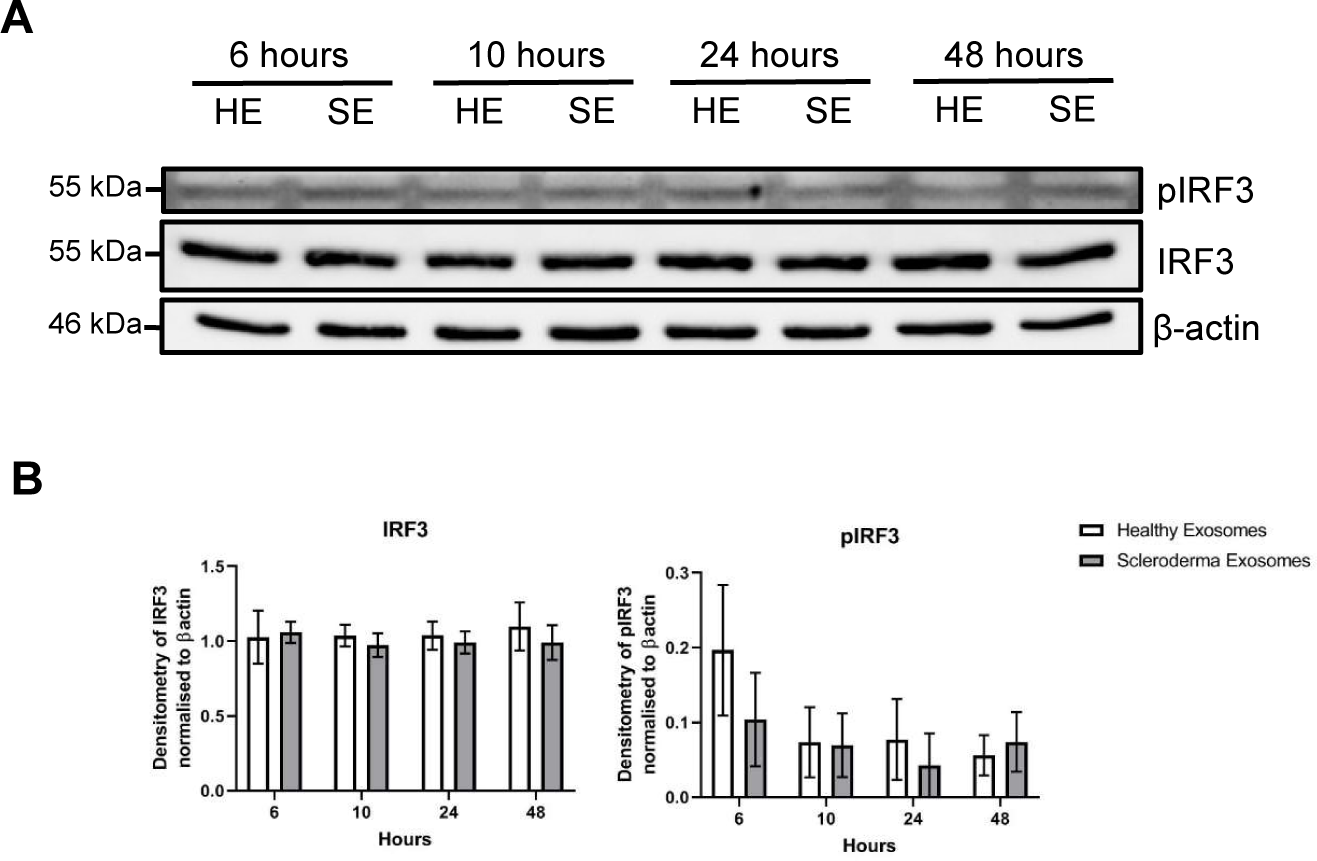
Scleroderma dermal fibroblast exosomes induce Type 1 IFN signalling in keratinocytes independent of IRF3/7. HaCat keratinocytes were stimulated with healthy and SSc fibroblast exosomes for 6-48 hours. Protein was isolated from stimulated HaCats. (A) pIRF3 and IRF3 protein levels were assessed by western blot. β-actin was used as a loading control. (B) Bar chart illustrates quantification of protein expression by densitometry and normalised to β-actin levels. Data represents means ± SE. Data was checked for normality and statistically analysed using a Two-way multiple comparison ANOVA, normalised by Tukey.

## Notes

### Competing Interest Statement

The authors have declared no competing interest.

