## Supplementary Materials and Methods for "Systemic Sclerosis dermal fibroblast exosomes trigger a Type 1 interferon response in keratinocytes through the TBK/JAK/STAT signalling axis"

**Patient cell lines**

Full thickness skin biopsies were surgically obtained from the forearms of four adult healthy controls and five adult patients with recent onset SSc, defined as a disease duration of less than 18 months from the appearance of clinically detectable skin induration. All patients satisfied the 2013 ACR/EULAR criteria for the classification of SSc. All participants provided written informed consent to participate in the study. Informed consent procedures were approved by NRES-011NE to FDG. Fibroblasts were isolated and established as previously described. SSc keratinocytes were grown from dissected skin biopsies and grown in keratinocyte grown media (Promega). Primary cells were immortalized using human telomerase reverse transcriptase (hTERT) to produce healthy control hTERT and SSc hTERT.

**Cell culture**

The immortalised keratinocyte cell line HaCats were maintained in Dulbecco’s modified Eagle medium (DMEM) (Gibco) supplemented with 10% FBS (Sigma) and penicillin-streptomycin (Sigma). HaCats were stimulated with 1% total volume of healthy and SSc fibroblasts exosomes for 48hours. TBK1 was inhibited with GSK8612 (10μM) and JAK1 inhibitor Tofacitinib (10μM) for 48hrs.

**Transwell Assay**

Fibroblasts were seeded onto 0.4-micron pore polyethylene terephthalate (PET) transmembranes (Corning). An equal number of HaCats were seeded into a separate cell-culture plate. Upon confluency, the pore was inserted into the well and incubated for 48 hours. The HaCats were harvested for RNA and protein analysis. The fibroblasts were stained with DAPI and counted to ensure equal amounts of fibroblasts were used in the experiment.

**Exosome Isolation**

Fibroblast cell cultures were grown as discussed above with exosome depleted FBS. Exosomes were isolated from media collected after 48-72hr by ultracentrifugation (42,000 rpm for 2 hours). The pelleted exosomes were isolated using the total exosome isolation reagent (Invitrogen) and re-suspended in PBS. RNA was then extracted and purified using the Total Exosome RNA & Protein Isolate Kit (Invitrogen), according to the manufactures protocol.

**Exosomal RNA transfections**

50ng of exosomal RNA isolated from healthy or SSc fibroblast exosomes was transfected into HaCats using lipofectamine 2000 (Invitrogen) and incubated for 48 hours. In addition, the exosome RNA was pre-treated with RNaseA for 2 hours at 37°C after which the RNaseA was inhibited with RNase inhibitor prior to transfection. Lipofectamine 2000 in the absence of RNA was added to the mock control cells.

**Immuno-labelling and visualisation of fibroblast-derived exosomes internalisation in human epidermal keratinocytes.**

Isolated fibroblast exosomes were stained Vybrant^TM^ DiO cell-labelling solution (Invitrogen) and added to HaCat culture media. Exosomes were added at 3, 6, 10, 24 and 48 hours, over a time-course. The HaCats were imaged on an LMS700 confocal scanning inverted microscope. The images were processed in Image J and Zen 3.1 (blue edition) software.

**CD63 ELISA**

Exosome abundance was measured using the ExoELISA-ULTRA CD63 enzyme-linked immunosorbent assay (ELISA) (System Biosciences). The ELISA plate is pre-coated with an anti-CD63 primary antibody, which recognises the tetraspanin CD63 on the exosome surface. The secondary antibody contains a horseradish peroxidase enzyme used for signal amplification. The colorimetric substrate 3,3’,5,5’-Tetramethylbenzidine (TMB) is used for the assay read-out. The results are read on a ThermoScientific Multiskan EX spectrophotometer at a wavelength of 450nm.

**Dynamic Light Scattering**

Exosomes were diluted in sterile PBS to a concentration of 100nM. The samples were injected (150μL, per sample) into a Wyatt miniDawnTreos® system (equipped with an additional DLS detector)..The raw data were analysed using the ASTRA 6.0.3® software supplied by the instrument, with the regularisation algorithm used to generate the size-distribution histograms. The histograms of signal intensity versus hydrodynamic radius were then plotted in OriginLab.

**Western Blotting**

Total proteins were extracted from fibroblasts in RIPA buffer and resolved by SDS-PAGE (10-15% Tris-Glycine). Proteins were transferred onto Hybond nitrocellulose membranes (Amersham biosciences) and probed with antibodies specific for pSTAT1, Total STAT1 (Cell signalling), CD63 (Abcam), TSG101(Abcam), Histone 3(Cell signalling), pIRF3 (Abcam), IRF3 (Cell signalling) and β-Actin (Sigma). Immunoblots were visualized with species-specific HRP conjugated secondary antibodies (Sigma) and ECL (Thermo/Pierce) on a Biorad chemiDoc imaging system.

**RNA sequencing**

Quality control of fastq files was performed with FastQC version 0.11.9. Reads were aligned to the human reference genome version GRCh38.84 using STAR aligner version 2.5.2b (21). MultiQC version 1.11 (22) was used to QC the alignment and generate an overall quality control report. Downstream analyses and visualizations were performed using R version 4.1.2 [R Core Team (2021). R: A language and environment for statistical computing. R Foundation for Statistical Computing, Vienna, Austria. URL https://www.R-project.org/]. BAM alignment files generated by STAR were transformed into count tables using Rsubread version 2.8.1 (23). The R package edgeR version 3.36.0 (24) was used for differential gene expression analysis. Gene set enrichment analysis for gene ontology biological processes was performed using WebGestalt (25, 26)

**RT-qPCR**

RNA was extracted from cells using commercial RNA extraction kits (Zymo Research). RNA (1ug) was reverse transcribed using cDNA synthesis kits (Thermo). QRT-PCRs were performed using SyBr Green PCR kits on a Thermocycler with primers specific for MX1 (Forward: CGACACGAGTTCCACAAATG Reverse: AAGCCTGGCAGCTCTCTACC), CXCL10 (Forward; GGTGAGAAGAGATGTCTGAATCC Reverse; GTCCATCCTTGGAAGCACTGCA), CXCL11 (Forward; TCCCCCATGTTCAAAAGAGGAC Reverse; ATATCTGCCACTTTCACTGCTTTTAC), OAS1 (Forward; CGGACCCTACAGGAAACTTG Reverse; GAAGCAGGAGGTCTCACCAG), IFIT1 (Forward; GACTGGCAGAAGCCCAGACT Reverse;GCGGAAGGGATTTGAAAGCT) and GAPDH (Forward; ACCCACTCCTCCACCTTTGA Reverse; CTGTTGCTGTAGCCAAATTCGT). Data were analysed using the ΔΔ Ct method. GAPDH served as a housekeeping gene. Type I IFN superarray was performed using manufacturers protocol (Qiagen)

**Transmission Electron Microscopy**

Healthy and SSc exosomes were loaded onto carbon-coated copper grids (Formvar/Carbon, 300 mesh Cu, Agar Scientific). Grids were glow discharged for 30 seconds at 0.39 mBar pressure and 10 mA (PELCO easiGlow, Ted Pella). Grids were incubated for 1 minute with 7 μL sample, washed three times with ddH_2_O and stained with 2% (w/v) uranyl acetate. Data were collected using a Tecnai F20 transmission electron microscope (FEI) at 200 KeV, fitted with a CETA CMOS CCD camera (FEI). Micrographs were collected at 40,000x magnification with a pixel size of 3.51 Å. Image J was used to add a scale bar and quantify the size of the vesicles.

**Immunohistochemistry**

Immunohistochemistry was performed as previously described (2). Sections were stained with a pSTAT1 (cell signalling 1/200), MX1 (abcam 1/100) and CXCL10 antibody (abcam 1/200), visualised using an HRP conjugated mouse secondary and counterstained with haematoxylin.

**Statistical Analysis**

Data are presented as the mean ± standard error. Statistical analysis was performed using a two-tailed, paired Student’s t-test for singular analysis and ANOVA tests for multi-comparison analysis.
